# UNCURL-App: Interactive database-driven analysis of scRNA-Seq data

**DOI:** 10.1101/2020.04.15.043737

**Authors:** Yue Zhang, Shunfu Mao, Sumit Mukherjee, Sreeram Kannan, Georg Seelig

## Abstract

**Motivation:** Analysis of single cell RNA sequencing (scRNA-seq) datasets is a complex and time-consuming process, requiring both biological knowledge and technical skill. With the rapid growth in scRNA-seq datasets, there is a need to simplify and systematize this process.

**Results:** We introduce UNCURL-App, an online GUI-based interactive tool which integrates the scRNA-seq analysis pipeline with prior knowledge. UNCURL-App introduces two key innovations: First, cell type databases are integrated directly with the rest of the analysis process, allowing the user to identify potential cell types or pathways directly from the data. Second, tools for interactive re-analysis allow the user to create, merge, or delete clusters. In addition, UNCURL-App integrates multiple stages of the data analysis pipeline into a single interface, including dimensionality reduction, clustering, differential expression, and cell type identification.

**Availability:** The web tool is available at https://uncurl.cs.washington.edu/. Source code is available at https://github.com/yjzhang/uncurl_app

## Introduction

Single cell RNA sequencing (scRNA-seq) has become an essential and ubiquitous tool for exploring the diversity of cell types in multicellular organisms. Progress in experimental technology development has driven rapid growth in the number of scRNA-seq datasets [1, 2], with a search in 2020 for ”single-cell RNA-seq” on NCBI GEO returning tens of thousands of results. Over little more than a decade, scRNA-seq experiments progressed from first proof-of-principle demonstrations using a handful of cells [3] to the construction of ”cell atlases” that enumerate all of the cell types present in an organ or organism [4–9]. However, while experimental approaches have become higher throughput and more widely available, it remains challenging to map experimentally determined single cell transcriptomes to biologically meaningful cell types. Given the very large throughput in cell number and the high complexity of many of the systems under investigation, reliable data analysis has become the main bottleneck of the scRNA-seq workflow.

The process of assigning sequenced cells to cell types is a multi-step process that requires the user to make decisions based on their judgment, because the ground truth about abundance and identity of cell types in an experiment of interest is typically not available. In practice, sequencing data is often first “pre-processed,” i.e. corrected for variability introduced by experimentally sampling the actual cellular transcriptome or “batch-corrected” if data from multiple experiments need to be integrated. Then, data are visualized in two dimensions and cells are clustered. Differential expression analysis identifies genes that are characteristic of each cluster, and these differentially expressed genes are used to assign clusters to cell types based on known gene-cell type associations. It is almost always necessary to iterate over this process and repeatedly remove, merge or split clusters to arrive at a satisfactory mapping of clusters to cell types consistent with known biology. These tasks require users who have both technical proficiency and knowledge of the underlying biology.

A wide range of computational tools have been developed to guide and assist each step in the analysis workflow from preprocessing and imputation [10–14], to clustering [15–19], data integration through batch effect correction [20, 21] and cell type annotation [22–24]. There are also a number of integrated analysis frameworks that combine several of these tasks into one package [25, 26]. However, these tools are typically restricted to command-line usage and require programming knowledge, hindering the accessibility of scRNA-seq analysis. These tools are also limited in their interactivity; even web-based tools such as scQuery [24] typically do not allow cluster assignments or cell type labels to be changed by the user. Moreover, in particular the last step of assigning labels to clusters remains heavily dependent on a user’s prior knowledge. Thus, even with all of these computational tools, the process of analyzing a new scRNA-seq data set remains somewhat idiosyncratic.

To aid in the task of analyzing scRNA-seq data, we here introduce UNCURL-App. UNCURL-App combines data preprocessing, dimensionality reduction, clustering, differential expression, and interactive data analysis within an online graphical user interface. UNCURL-App introduces two key innovations: First, cell type databases are integrated directly with the rest of the analysis process, accelerating mapping of clusters to cell types. Second, UNCURL-App includes tools for interactive re-analysis that allow the user to create, merge, or delete clusters, thus making it possible to iteratively refine clusters using knowledge about gene expression, putative cell type annotations, and other information accessible through UNCURL-App. Because the entire workflow can be performed in a browser and because external knowledge is made available during the analysis process, we expect UNCURL-App not only to accelerate scRNA-seq analysis but also to further extend the user base for this technology.

## Materials and methods

### Workflow & Interface

From the user’s perspective, the first step in the UNCURL-App pipeline is to upload the data as a gene-cell read count matrix (Supplemental Figures 1 & 2). After uploading the data, the raw gene-cell matrix goes through a few basic transformations including data filtering and normalization before being used as input into UNCURL [13]. These steps are described in the Supplementary Methods. Next, the app performs data decomposition using UNCURL, dimensionality reduction, and differential expression, which is detailed below. Then, the user is redirected to an interactive web site, from which they can view the results of the previous steps, query for cell types, or perform interactive re-analysis (Figure 1).

**Fig 1.**
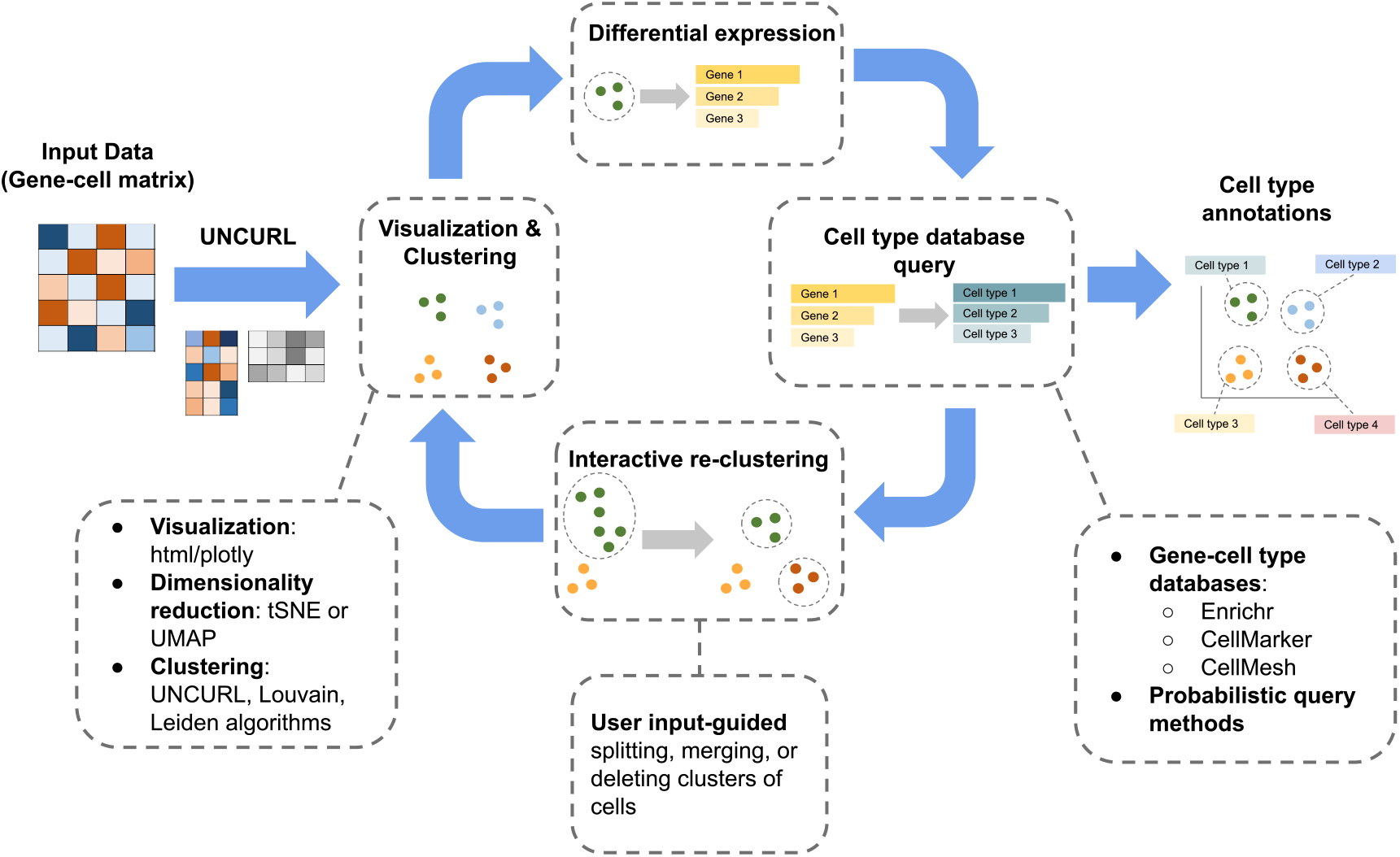
This shows the major components and workflow of UNCURL-App. Given the user-uploaded gene-cell matrix: 1) UNCURL is run on the data, producing a clustering; 2) dimensionality reduction is used for visualization, and 3) differentially expressed genes are identified. Then, the user may do cell type annotation using database queries on the top genes, and perform interactive re-clustering.

There are three main visualization components: the dimensionality-reduced scatterplot of cells, the barplot showing the most differentially expressed genes, and the cell type query results. A labeled screenshot of the main UNCURL-App view is shown in Figure 2. The scatterplot, on the top left of the screen, shows a dimensionality-reduced view of the cells in the dataset, where each point represents a cell. This view can be colored by cluster, gene expression for selected gene(s), or custom label sets based on uploaded files or user-defined criteria. For example, a user may select all cells belonging to a given cluster that also have positive expression of a certain gene, or select all cells that both belong to a certain cluster and have a certain label in an uploaded file (Supplemental Figure 3). The user may also select cells by drawing a box or shape on the plot itself.

**Fig 2.**
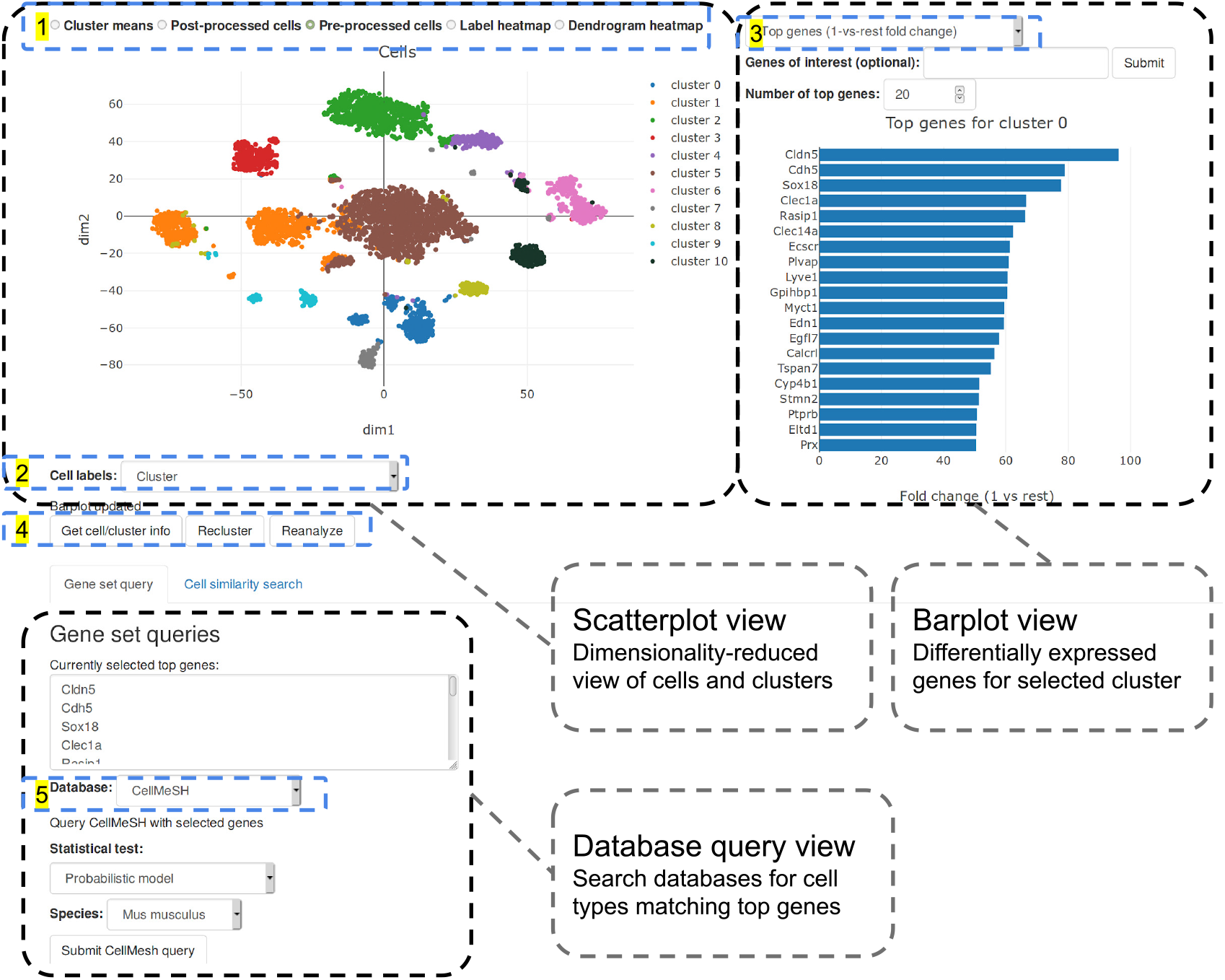
This figure shows the main user interface of UNCURL-App. The top left contains the scatterplot, showing a views of the cells and clusters. The top right is a barplot containing the top genes per cluster. Clicking on a cluster in the scatterplot updates the top genes. On the bottom is the database query view, which is used to identify cell types present in the dataset. **Options: (1)** These are options for what type of visualization to show in the scatterplot.”Cluster means” shows one point for each cluster. ”Post-processed cells” shows a dimensionality-reduced view of cells that takes the *W* matrix returned by UNCURL as input. ”Pre-processed cells” (default) shows a similar view, but using the original data as input for dimensionality reduction. ”Label heatmap” shows a comparison of two cell label sets. ”Dendrogram heatmap” shows a clusters vs genes dendrogram. **(2)** Options for how to color the cells/points in the scatterplot. Can be based on cluster, gene expression, or a custom label set. **(3)** Options for how to identify the top genes. Can be 1-vs-rest or pairwise, p-value or ratio. **(4)** ”Get cell/cluster info”: get read/gene count for the selected cell and cluster. ”Recluster”: options to merge, split, delete, or create a new cluster. ”Reanalyze”: options to re-run the whole analysis. **(5)** There are a number of databases that can be used for cell type identification, including CellMarker, CellMeSH, and Gene Ontology.

On the top right of the screen, the barplot shows the top differentially expressed genes for the selected cluster or label. The barplot is automatically updated whenever the user clicks on a cell on the scatterplot, showing the top genes for the cluster that the cell belongs to.

The bottom left of the screen shows the database query view. From this view, the user may query cell type databases using the top differentially expressed genes. Databases include Enrichr [27], CellMarker [28], and Gene Ontology [29, 30], as well as our new cell type database, CellMeSH [31]. Submitting a query will return a list of cell types with a confidence score, overlapping genes, and references for each gene-cell type pair.

A help page for UNCURL-App detailing the interface and available options is included in the web app, and can be accessed at https://uncurl.cs.washington.edu/help.

### Data preprocessing, clustering, and differential expression

The first step in the analysis pipeline builds on UNCURL, a tool for data decomposition and clustering scRNA-seq data using probabilistic matrix factorization [13]. UNCURL has been shown to have state-of-the-art performance in clustering large-scale scRNA-seq datasets, and performs exceptionally well on sparse datasets. It assumes that the observed read count matrix is distributed with either a Poisson, Log-Normal, or Gaussian distribution, with the parameters of the distribution coming from a hidden state matrix. This hidden matrix is the product of two non-negative matrices of rank *k: M*, the archetype matrix, of shape *genes × k*, and *W*, the weights matrix, of shape *k* × *cells*, where each column sums to 1. These two matrices are the outputs of UNCURL. The rank *k* can be manually set as an input parameter, or automatically determined using the gap score [32] if *k* is set to 0. By default, *k* is set to 10, which tends to produce good results in practice, but can be changed interactively by merging or splitting clusters as discussed in more detail below.

The result of UNCURL is then used for dimensionality reduction and clustering. Dimensionality reduction is done using standard methods, such as tSNE [33], UMAP [34], or PCA. This produces a two-dimensional scatterplot of cells. By default, clustering is done using argmax on the *W* matrix returned by UNCURL (as described in [13]). Each column in *W* represents the weights for each archetype in one cell, so the archetype with the maximum weight is the most likely cluster assignment for that cell. Clustering can also be done using the Louvain [35] or Leiden [36] community detection algorithms, which also use the *W* matrix as input. However, only clustering using UNCURL is compatible with iterative cluster refinement as detailed below.

In order to identify the most differentially expressed genes in each cluster, UNCURL-App uses one of two methods: the t-test, or the ratio of means. These metrics can either be calculated for one cluster against all other clusters, or against a single cluster. The t-test has been shown to be one of the best performing methods for identifying DE genes in scRNA-seq datasets, and is also much faster than more complex methods [37].

### Interactive data analysis

UNCURL-App has the capacity to merge, split, or delete clusters of cells in an interactive fashion. After the initial analysis process is completed, there are often refinements to the clustering that users would like to make, no matter the quality of the initial clustering. For example, the user may want to split a large cluster, merge multiple similar clusters, or delete a group of poor quality cells or potential doublets. This cluster refinement might be based on the shape of the scatterplot, differential expression results, cell type queries, or some other metrics.

The user-driven changes in clustering are incorporated into UNCURL by using them to generate new initializations and then re-running the optimization process, as shown in Figure 3. This process fundamentally relies on the UNCURL algorithm [13], and was inspired by the UTOPIAN software for interactive non-negative matrix factorization [38], but in UNCURL-App, cells take the place of documents. Say that we have matrices *M* and *W*, with shapes *g × K* and *K × c*. In order to split a selected cluster, we first run k-means with *k* = 2 on the cells assigned to the selected cluster. This generates new matrices *M_cluster_* and *W_cluster_*, of shape *g* × 2 and 2 × *c* representing the means and cell cluster assignments. The column and row corresponding to the selected cluster are deleted from *M* and *W*, and *M_cluster_* and *W_cluster_* are appended to *M* and *W*, creating *M_new_* and *W_new_*, with shapes *g × (K* + 1) and (*K* +1) × *c*. Then, UNCURL is re-run with *M_new_* and *W_new_* as the initializations, which affects other clusters as well. The process is analogous for merging clusters and assigning cells to new clusters: we create new initializations for *M* and *W* using the selected clusters or cells, and then re-run the optimization process.

**Fig 3.**
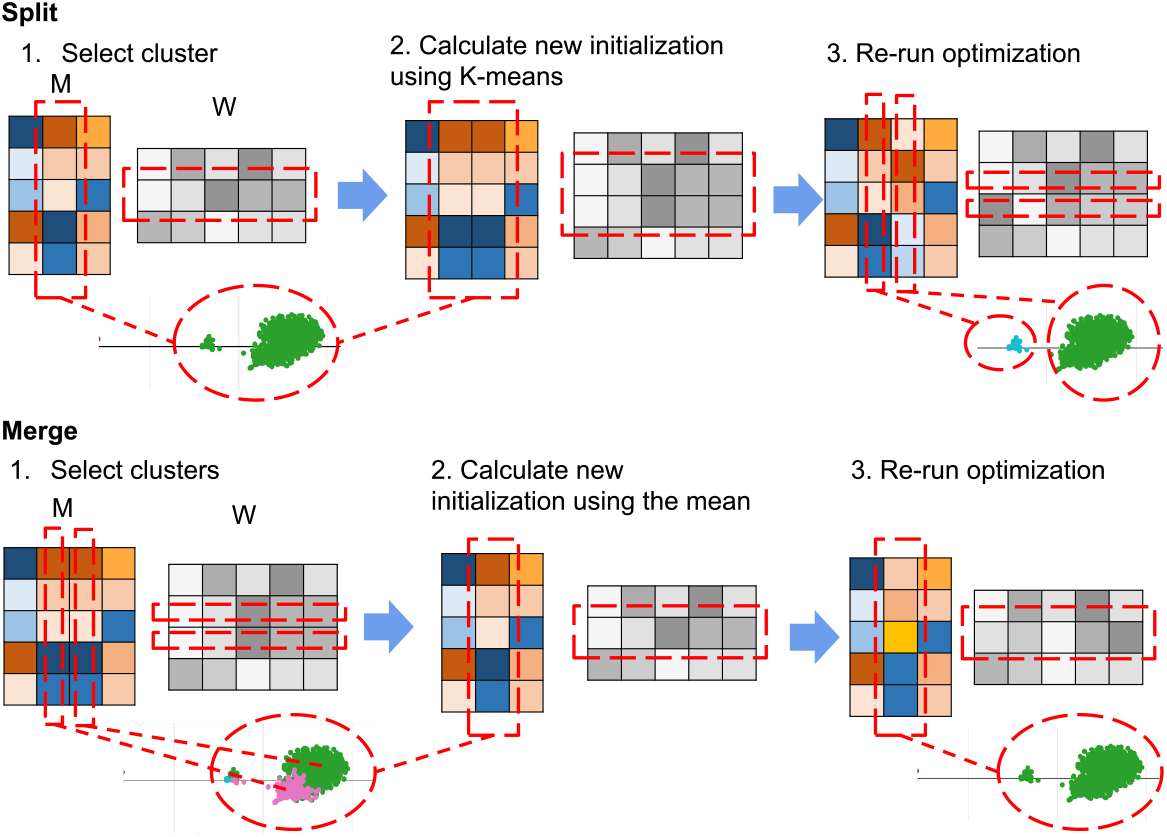
Splitting/Merging clusters. This shows how the process of splitting and merging clusters works in UNCURL-App. Both processes begin with the *M* and *W* matrices returned by UNCURL, and the cluster(s) to merge or split. In order to split a cluster, a new initialization for *M* and *W* is created by running k-means on the cluster of cells to be split. Then, the UNCURL optimization process is re-run to create new matrices. The process for merging is analogous: a new initialization is created using the mean of the selected cells, and then the UNCURL optimization process is re-run to create a new *M* and *W*.

After generating *M_new_* and *W_new_*, a new visualization, clustering, and differential expression results are calculated using *W_new_*. Running re-clustering also automatically updates the differential expression results. In addition, UNCURL-App saves a history snapshot whenever one of the re-clustering operations is run, allowing the user to restore the state prior to changing the clustering.

### Cell type annotation

UNCURL-App has a number of interfaces to external databases, which are used to assist with identifying cell types present in the dataset, as well as helping to better understand underlying biological processes. First, UNCURL-App contains an interface to the Enrichr tool for gene set analysis [27, 39]. This tool contains interfaces to a variety of gene set databases that can be used to help identify cell function. We also provide an interface to Gene Ontology [29, 30], which is queried using the goatools package [40]. In addition, we have two databases specifically for cell type identification, CellMarker and CellMeSH.

CellMarker is a hand-curated database of cell types, annotated with marker genes based on a literature search [28]. This dataset consists of 673 cell types, where each cell type is associated with an average of 72 and a median of 9 marker genes. To search this database given a list of query genes, we use the hypergeometric test for the overlap between the query gene set and the marker genes.

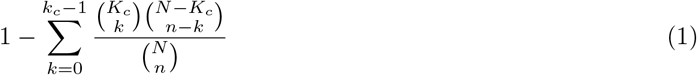

where *N* is the total number of genes, *n* is the number of genes in the query set, *K_c_* is the number of genes for the cell type, and *k_c_* is the number of genes that overlap between the query and the cell type. This is the probability that, given that the query gene set is randomly sampled, the overlap is greater than or equal to the actual overlap. To find the top cell types for a query gene set, this p-value is calculated for all cell types and ranked in ascending order.

CellMeSH is a new database that maps cell types to their associated genes [31]. It was created by combining two existing literature indices: the MEDLINE citation index [41], which contains publication abstracts with associated metadata, and the gene2pubmed database [42], which contains a mapping of genes to publications. The key metadata from MEDLINE are the associated Medical Subject Headings, or MeSH terms [43], a subset of which represent cell types. For each cell type from MeSH, we found all publications where they occur, and all genes that occur in the same publications, thus creating an association between cell types and genes. This database contains 570 cell types, 292 of which with at least one associated gene. Searching this database can be done using a hypergeometric test or the probabilistic method described in [31].

## Results

### Examples

In order to validate the UNCURL-App workflow, we used the app to analyze three different scRNA-seq datasets, as described below. For these datasets, we performed clustering and cell type annotation using UNCURL-App with default settings. Cell type labels were generated by querying the top 50 genes by 1-vs-rest ratio with the CellMeSH database (see Methods).

The running times of the non-interactive steps are shown in Table 1. The running time for UNCURL scales linearly with the number of cells, while the running time for tSNE scales with order *n* log *n*, where *n* is the number of cells. With larger numbers of cells, running tSNE is the most time-consuming step. This can be obviated by using UMAP as the dimensionality reduction method.

**Table 1.**
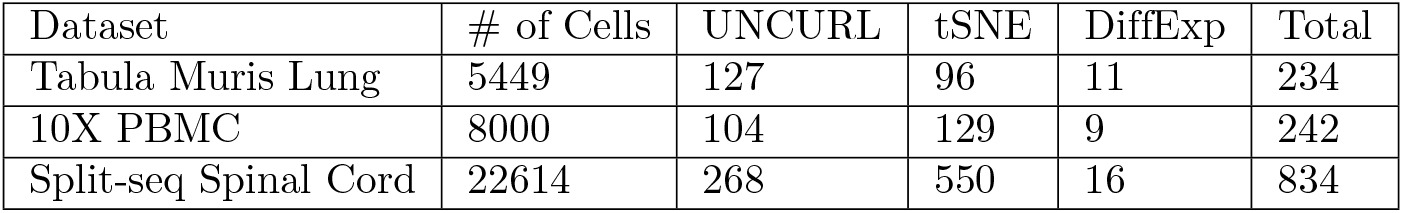
Runtime of UNCURL-App. These times were based on an Amazon Web Services (AWS) t2.medium instance, with two processors and 4GB of memory. All times are in seconds.

#### Example: Tabula Muris lung cells

As a first example, we consider a subset of the Tabula Muris dataset from [7] containing only cell types found in the lung. This dataset contains 5449 cells and 14 annotated cell types. The labels in the original study were generated by first running graph-based clustering and then manually examining the marker genes for each cluster.

After uploading the dataset and processing it with default settings, we see the clustering and initial cell type assignments in Figure 4a. The clustering was based on UNCURL, and the scatterplot visualization was generated using tSNE. Based on the scatterplot, cluster 4 appeared to consist of at least two distinct groups of cells. In addition, the top cell types from a CellMeSH query on the top genes in this cluster included both B and T cells (Supplementary Figure 4a), suggesting that this cluster might be a mixture of at least these two cell types. Based on these observations, we decided to split this cluster using our interactive data analysis tools, resulting in the clusters given in Figure 4b. The post-split cell type assignments (Figure 4c) appeared to be more consistent with known biology than the original assignments.

**Fig 4.**
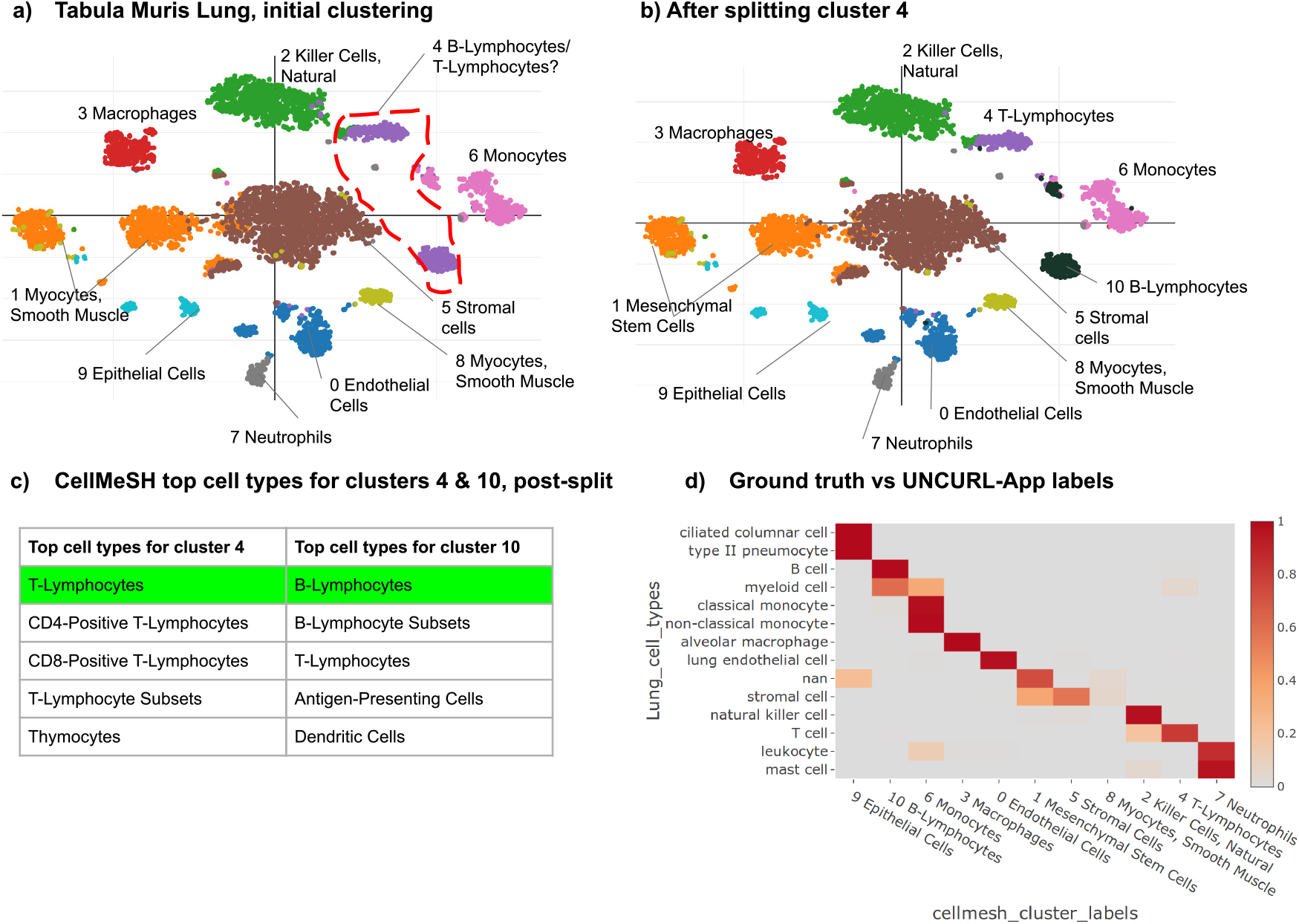
Tabula Muris Lung. Examples of running UNCURL-App on the Tabula Muris lung dataset **(a) (b)** These show the same tSNE scatterplot, with clusters from the initial clustering and labels from CellMeSH and the clusters and labels after splitting cluster 4 **(c)** These show the most relevant cell types returned by CellMeSH for the given clusters. Cell types matching ground truth labels are highlighted in green. **(d)** This is a heatmap showing the relationship between the ground truth clusters and the clusters generated by UNCURL-App, after the split. The colors indicate the proportion of the ground truth cluster (left) that overlap with the UNCURL-App cluster (bottom). The final NMI (normalized mutual information score) between the published labels and the inferred clusters is 0.80.

Based on Figure 4d, there is generally good concordance between the generated clusters and original clusters, as well as between assigned labels and the original labels. Of the labels that were different, in most cases UNCURL-App assigned cell types that were closely related to the original ground truth label (for example, pneumocytes and columnar cells are subsets of epithelial cells, and neutrophils are a subset of leukocytes). The stromal cells are split into multiple clusters in UNCURL-App, which could represent heterogeneity in the original sample that was not captured by the original labels. No prior information about the cell types present in the dataset was used at any point in this process.

#### Example: 10X PBMCs

Next, we turned to a dataset comprised of 8000 human peripheral blood mononuclear cells (PBMCs) from [44]. This dataset was created by randomly sampling 1000 cells from each of 8 scRNA-seq datasets comprised of cells that were flow-sorted based on known cell-type markers. Thus, the ground truth cell type labels represent pure samples, as opposed to the computational assignments used as ground truth in the other example datasets.

UNCURL-App was run with default settings to generate 10 initial clusters (Figure 5a). Looking at the resulting clusters and putative cell type assignments (5b), it appeared that clusters 2 and 6, labeled Neutrophils and Monocytes, were very similar, and could just represent a single group of cells. A pairwise differential expression analysis (Figure 5c) further illustrates that only related genes, S100A8 and S100A9, appear to be significantly differentially expressed between these clusters. Plotting the expression levels of these genes (Figure 5d), it seems that the small group of cells to the left of the main cluster has much higher expression of these genes, suggesting that this group might constitute a separate cluster. Thus, we first merged clusters 2 and 6, and then split off that small group of cells. These operations resulted in the clustering shown in Figure 5e.

**Fig 5.**
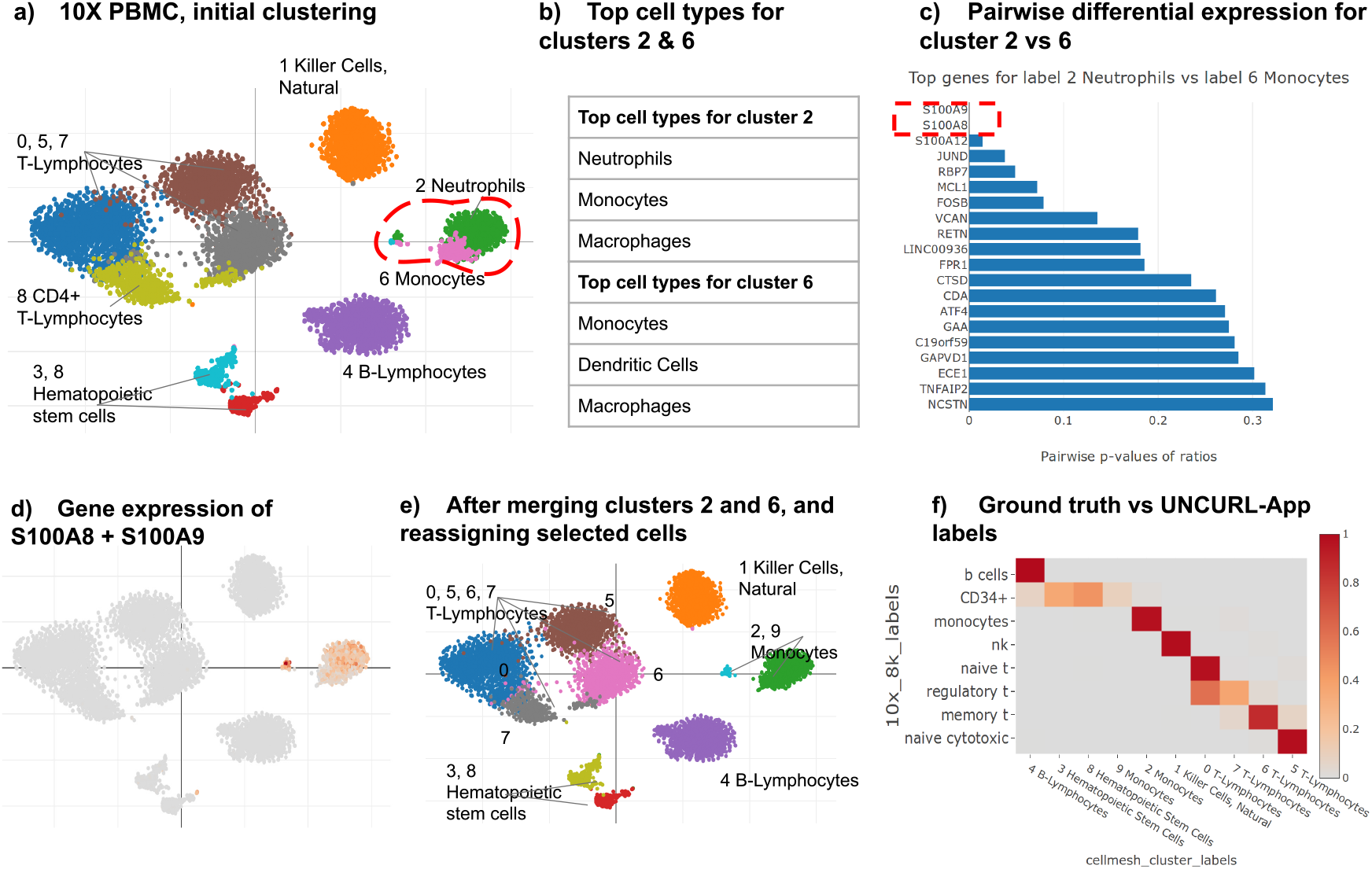
10X PBMC. Examples of running UNCURL-App on the 10X PBMC dataset **(a) (e)** These show the same tSNE scatterplot, with clusters from the initial clustering and labels from CellMeSH, and the clusters and labels after merging clusters 2 and 6 and creating cluster 9,. **(b)** This shows the most relevant cell types returned by CellMeSH for the given clusters. **(c)** This shows the top differentially expressed genes for Cluster 2 vs Cluster 6. **(d)** This shows the sum of the gene expressions of the two genes S100A8 and S100A9. **(f)** This is a heatmap showing the relationship between the ground truth clusters and the clusters generated by UNCURL-App, after the split. The final NMI (normalized mutual information score) between the true labels and the inferred clusters is 0.84.

As with the previous dataset, there now is good correspondence with the ground truth clusters and labels (Figure 5f). Cells of the same ground truth type are generally assigned to the same cluster, and the cluster labels returned by CellMeSH generally correspond to the ground truth labels. CD34+ cells are generally recognized as hematopoietic stem cells [45], so the CellMeSH label here seems to be accurate. One major difference is that CellMeSH labeled all four T cell subtypes as ”T-Lymphocytes”, even though they were clustered into distinct clusters. To investigate further, we looked at the full list of CellMeSH labels for these clusters, not just the top one. These results are shown in Supplementary Figure 5b, with the cell types most similar to the ground truth highlighted in green. For example, Cluster 0 corresponds to naive T-cells, which are selected as CD4+. Cluster 5 corresponds to naive cytotoxic T-cells, which are CD8+, and the ”CD8+ T-Lymphocytes” label is the third highest label, below ”T-Lymphocytes” and ”Lymphocytes”. Cluster 6 corresponds to memory T-cells, which can be either CD8+ or CD4+; the second and third labels are ”CD8+ T-Lymphocytes” and ”CD4-Positive T-Lymphocytes”. Cluster 7 corresponds largely to regulatory T-cells, which are CD4+, and the second and third highest CellMeSH labels are ”CD4-Positive T-Lymphocytes” and ”T-Lymphocytes, Regulatory”. This shows a good correspondence between the true and assigned labels at a more fine-grained level.

#### Example: SPLiT-seq spinal cord

For a final test we turned to a larger dataset comprised of 22,614 mouse spinal cord nuclei from 2 and 11-day old mice sequenced using SPLiT-seq [9]. This dataset has 44 annotated cell types, which is substantially more than the previous two datasets. However, many of these annotated cell types are closely related (for example, there are 15 types of excitatory neurons), so for the ”ground truth” comparisons in this section, we combine many of the annotated cell types into larger clusters of similar cells. Even after this process, many of the cell types are similar, with many subtypes of neurons.

We first ran UNCURL-App with default settings to generate an initial clustering with 10 clusters, several of which exhibit substantial heterogeneity (Figure 6a). For example, cluster 8 (labeled as ”Endothelial Cells”) represents at least four different groups of non-neuronal cells. Thus, we split them into four different clusters (Figure 6b). It is clear that splitting the clusters worked to separate what appeared to be distinct cell types. In addition, the clusters that CellMeSH labels as ”Neurons” or ”Interneurons” (3, 6, 9, 0) all appear to be rather heterogeneous. Results after splitting some of the neuronal clusters are shown in Supplemental Figures 6-9.

**Fig 6.**
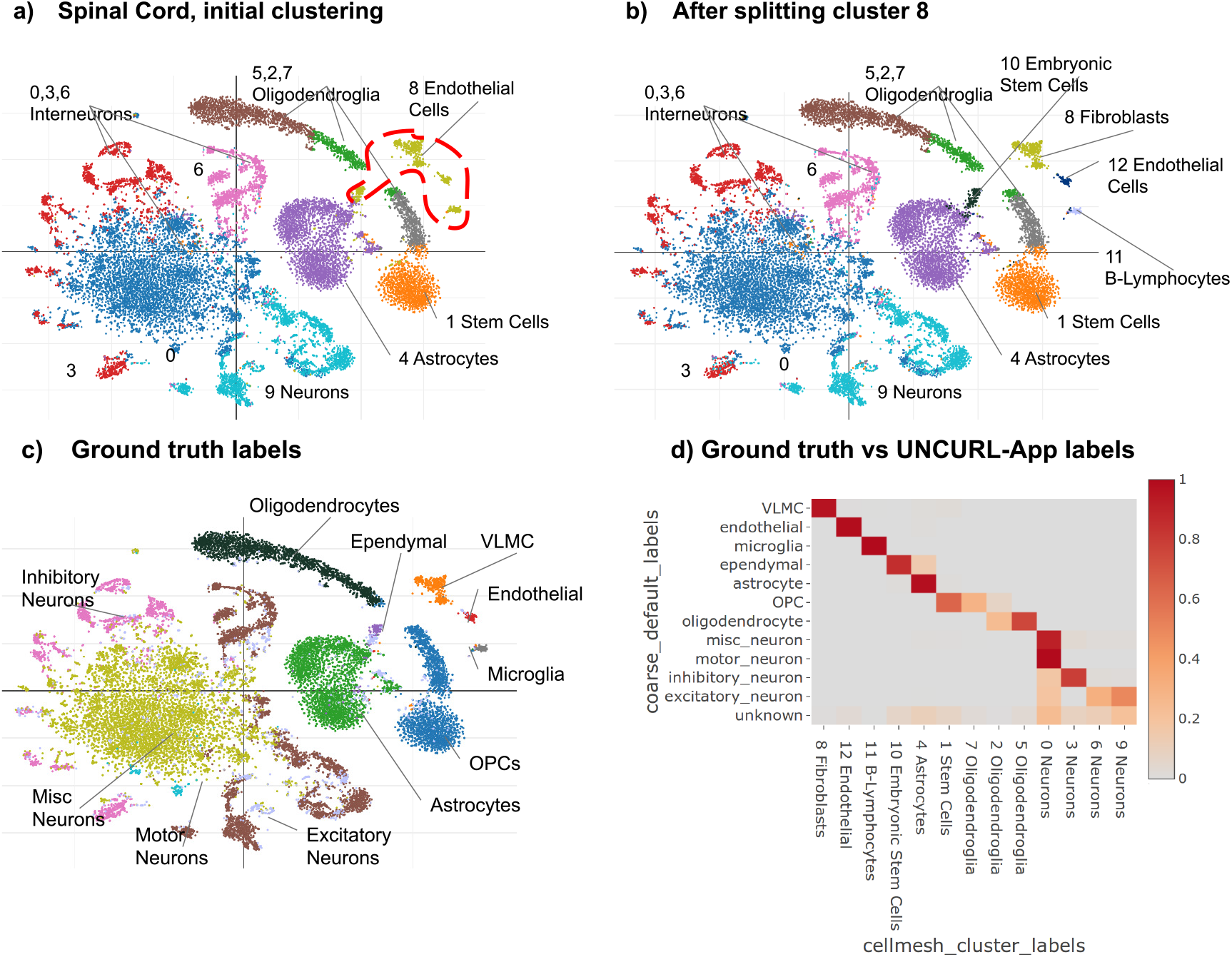
Spinal Cord. Examples of running UNCURL-App on the split-seq spinal cord dataset **(a)** tSNE scatterplot, with 10 clusters from the initial clustering and labels from CellMeSH. Cluster 8 which consists of multiple disconnected groups of cells is highlighted. **(b)** Clusters and labels from CellMeSH after splitting cluster 8. **(c)** Clusters and labels after refinement with ground truth labels from the original dataset. **(d)** This is a heatmap showing the relationship between the ground truth clusters and the clusters generated by UNCURL-App, after the split. The final NMI (normalized mutual information score) between the published labels and the inferred clusters is 0.68.

As with the previous datasets, there is generally good concordance between the cell types from the original paper and the clusters generated by UNCURL-App, as shown in Figure 6d. Also similarly to previous datasets, the CellMeSH annotations were generally coarser grained than the original hand-annotated labels, with all of the neuron clusters being labeled as ”Interneurons” or just ”Neurons”. For the non-neuronal results, interpreting the labels identified by CellMeSH is more challenging (Supplemental Figure 7). Oligodendrocytes, astrocytes, and endothelial cells were correctly identified. For cluster 8, the ground-truth label was ”VLMC”, or ”vascular and leptomeningeal cells”. This is a highly specific category that does not appear in the CellMeSH ontology but was used as a cell label in [46]. Still, while coarse, the first three labels suggested by CellMeSH (Stromal Cells, Fibroblasts, Mesenchymal Stem Cells) seem consistent with cells derived from the meninges, the membrane enveloping the brain and spinal cord. In cluster 10, the ground-truth label ”Ependymal” was not correctly identified by CellMeSH, and the returned results did not seem to relate to ependymal cells. This points to a paucity of annotated publications with gene markers for this cell type. For cluster 11, all of the top CellMeSH results were immune cells, a group which the published label, ”microglia”, belongs to. ”Microglia” was one of the top 10 cell types returned.

## Discussion

### Dealing with unknown cell types

In many circumstances, scRNA-seq experiments will reveal cell populations that are distinct from any known cell types. This is a notable advantage of scRNA-seq, but it makes the problem of cell type annotation somewhat ill-defined. With UNCURL-App, if a user attempts to annotate a previously unknown cell type with CellMeSH, then CellMeSH will return the closest cell type present in the database. Even if the exact cell type is not present, this still provides useful information, as the returned cell type is usually a supertype of the more specific cell type of interest. In addition, UNCURL-App allows for gene set queries to other databases, such as the Gene Ontology [29, 30] and KEGG Pathway databases [47], that can be used to aid in cell type identification. This might be a supertype of the more specific cell type of interest.

### Comparison with existing tools

Unlike other general-purpose toolkits for scRNA-seq analysis such as scanpy [25], Seurat [26], and Monocle 2 [48], UNCURL-App is a web-based GUI tool that does not require command line usage. This allows a much wider range of potential users, such as biologists who are not programmers.

There are a number of web-based tools for scRNA-seq analysis, such as scQuery [24], Granatum [49], Cumulus [50], and Galaxy [51]. All of these tools, along with UNCURl-App, can perform dimensionality reduction, clustering, and differential expression on uploaded single-cell datasets. Both scQuery and Cumulus (along with UNCURL-App) but not Granatum or Galaxy also do cell type annotation. Cumulus requires users to provide cell type gene markers in order to annotate cell types, while UNCURL-App does not require any user input for cell type annotation, as it uses the CellMeSH database. ScQuery identifies cell types using cell type annotations derived from GEO. Compared to scQuery, the annotation accuracy of CellMeSH (used by UNCURL-App) is significantly higher [31]. Unlike all of the aforementioned tools, UNCURL-App is also capable of interactively merging, splitting, and deleting clusters of cells within the GUI.

There are a number of tools that classify cells given gene markers for known cell types, such as [22, 52]. We view these tools as complementary to UNCURL-App and CellMeSH. These tools require some knowledge of the cell types present in the dataset, as well as a way to manually find gene markers for these cell types, whereas such prior knowledge is unnecessary in the UNCURL-App/CellMeSH pipeline. In addition, CellMeSH can be used to improve the workflow for these tools by automatically selecting gene markers, obviating the need for manually finding them.

There also exist tools that perform single cell similarity search on reference datasets, such as CellAtlasSearch and scMatch [23, 53]. Rather than using marker genes, these methods compare the entire gene expression profile of every single cell to a reference database, using locality-sensitive hashing in the case of CellAtasSearch [23] or Pearson or Spearman correlation in the case of scMatch [53]. These tools do not include functionality for clustering or low-dimensional visualization. The advantage of UNCURL-App comes with its integration of clustering, differential expression, interactive re-analysis, and cell type querying into one easy-to-use platform.

## Conclusion

UNCURL-App provides a useful way to perform interactive scRNA-seq data analysis, including cell type annotation. In the future, we hope to augment UNCURL-App with new analysis capabilities, such as cell lineage and gene network analysis. We also hope to connect UNCURL-App to additional sources of information for cell type and functional annotation. This could come in the form of connections to new databases, or expansions to the CellMeSH database. Our ultimate goal is to increase UNCURL-App’s utility as a general tool for scRNA-seq analysis.

## Acknowledgements

We thanks Li Liu, Tao Peng, and Matthew Hirano for testing UNCURL-App and suggesting new features.

