## supplement for "UNCURL-App: Interactive database-driven analysis of scRNA-Seq data"

### Supplementary Information for UNCURL-App: Interactive Database-Driven Analysis of scRNA-Seq Data

#### 1 Implementation

UNCURL-App and the associated backend tools and databases are written in Python. The primary package is the uncurl-app package, which uses the Flask library as the server backend. Visualization is done in javascript using the plotly library. The backend, which interfaces with the dimensionality reduction and differential expression methods, is provided by the uncurl-analysis package, and the databases are provided by the cellmarker and cellmesh packages.

##### 1.1 Deployment

UNCURL-App has been tested to run on Ubuntu 16.04 and above, and can be deployed on a local or cloud server using Docker. We have created an example UNCURL-App deployment at https://uncurl.cs.washington.edu/. This deployment limits its upload size to 150MB.

#### 2 Pre-processing Details

UNCURl-App has a number of options for pre-processing the data prior to running UNCURL, which are described below.

##### 2.1 Batch Effect Correction

When running UNCURL-App with multiple datasets, there is the option of using batch effect correction to integrate the data. This uses the mutual nearest neighbors approach [1], as implemented in mnnpy [2].

##### 2.2 Data Filtering

By default, UNCURL-App removes the top and bottom 5% of cells by total gene expression level. This threshold can be adjusted.

Genes are sub-selected based on variance: first, the genes are binned into $n$ bins based on mean gene expression level. Next, for each bin, the top $k$-fraction of genes are selected based on variance. The default values for $n$ and $k$ are 5 and 0.2. This is the same process that is used in [3].

#### 2.3 Data Normalization

By default, UNCURL-App normalizes the data by dividing the value of every gene in each cell by the sum of gene expression levels in that cell, so that the total gene expression level sums to 1, and then multiplying the expression levels by the median gene expression levels across all cells. This is the same process that is used in [3]. This is done before gene selection. The normalized and filtered data is then used for running UNCURL, as well as for differential expression analysis.

### 3 User Interface

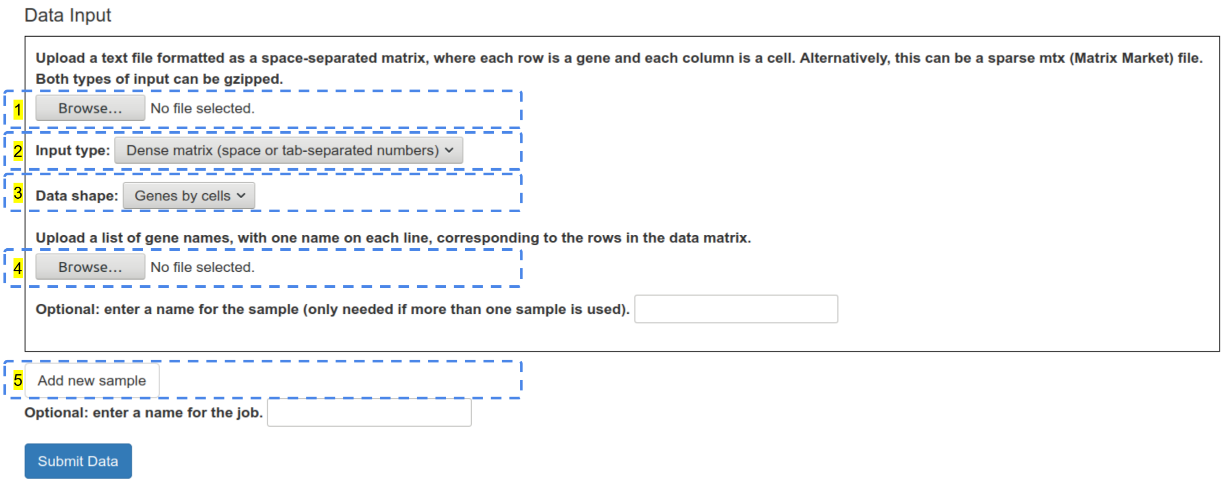


Figure 1: Data input interface of UNCURL-App. **(1, 2)** The scRNA-seq data uploaded can be a tab-separated file, or a sparse matrix in the .mtx format. **(3)** The input matrix shape can be genes by cells or cells by genes. **(4)** A separate text file containing gene names should be uploaded, with one gene name per line. **(5)** UNCURL-App can be used with multiple samples; this allows the user to upload additional data matrices.

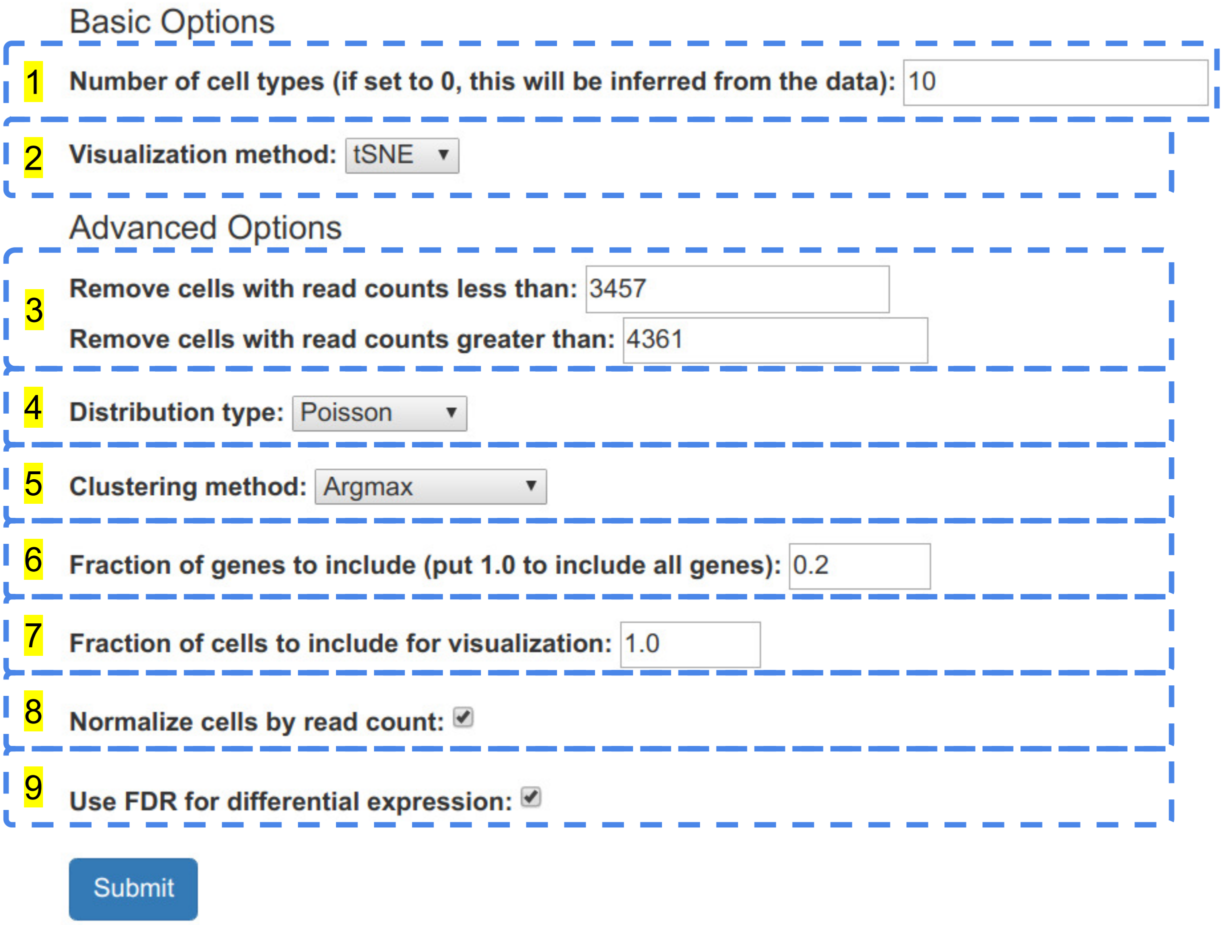


Figure 2: View after uploading a dataset and options that can be used for analysis. **(1)** The number of initial cell types (default: 10); if set to 0, this is automatically inferred using the gap score metric. **(2)** The dimensionality reduction method used for visualization can be tSNE, PCA, or UMAP. **(3)** Option to filter cells by min/max read counts. By default, this is set to the bottom and top 5%. **(4)** The distribution type is an option used by UNCURL, representing the sampling distribution of the dataset. This is Poisson by default, but could also be Log-Normal or Gaussian. **(5)** The default clustering method is Argmax with Louvain or Leiden as alternative options. **(6)** By default, UNCURL selects genes by first binning the genes into 5 bins by mean, and selecting the top 20% of genes by variance within each bin. The fraction can be set to any number between 0.01 and 1. **(7)** If this parameter is less than 1, then a random sample of cells is selected for visualization. **(8)** If this option is checked, then before running any other analysis step, the input matrix is normalized so that the total read count per cell is approximately the same. **(9)** If this is checked, the p-values shown for the differential expression will be adjusted using the Benjamini-Hochberg procedure for false discovery rate.

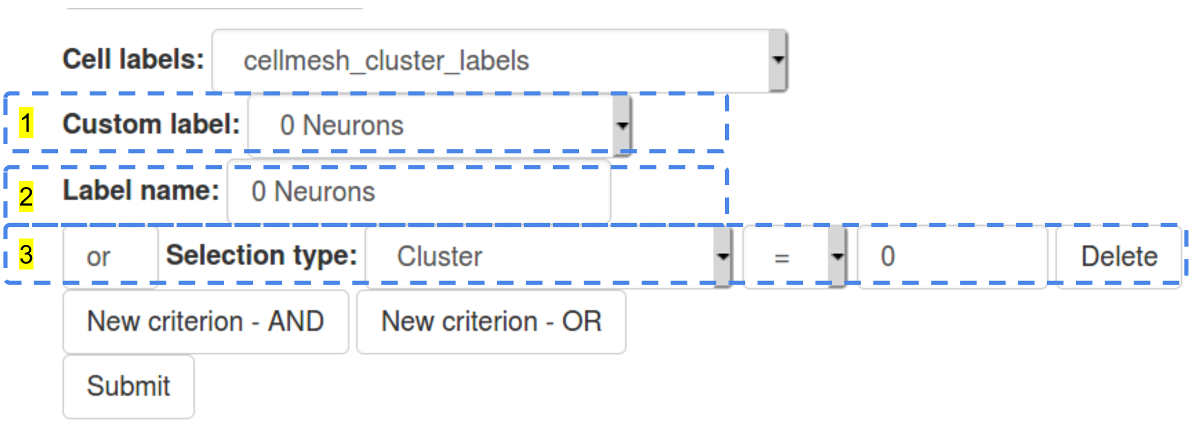


Figure 3: User interface for creating custom cell selections. Custom cell selections provide a way to create labels based on user-defined criteria. This view is shown after selecting ”Custom cell selection” from the ”Cell labels” dropdown menu. After creating a name for the cell selection (in this case, ”cellmesh_cluster_labels”), the user can create custom groups of cells using a variety of criteria. **(1)** Each ”Custom label” is essentially a cluster of cells defined using custom criteria. **(2)** Input for naming the custom label. **(3)** For each custom label, there can be any number of selection criteria. The ”Selection type” may be UNCURL-App clusters, user-uploaded labels, gene expression values, or selections drawn on the scatterplot using Plotly's box or lasso select tools. New selection criteria can be added using the ”New criterion - AND” and ”New criterion - OR” buttons below.

### 4 Tabula Muris dataset

**a) CellMeSH top cell types for cluster 4**

| Top cell types for cluster 4 |
|---|
| B-Lymphocytes |
| T-Lymphocytes |
| B-Lymphocyte Subsets |
| T-Lymphocyte Subsets |
| T-Lymphocytes, Helper-Inducer |

**b) Ground truth labels**

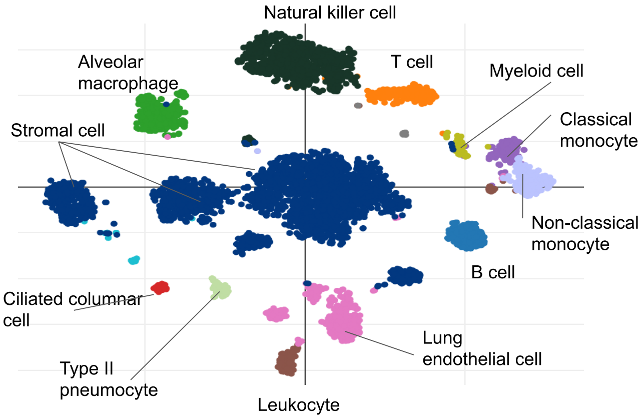


Figure 4: **a)** Top cell types for cluster 4 (in Figure 4a) **b)** tSNE plot with ground truth labels

### 5 10X PBMC dataset

**a) Ground truth labels**

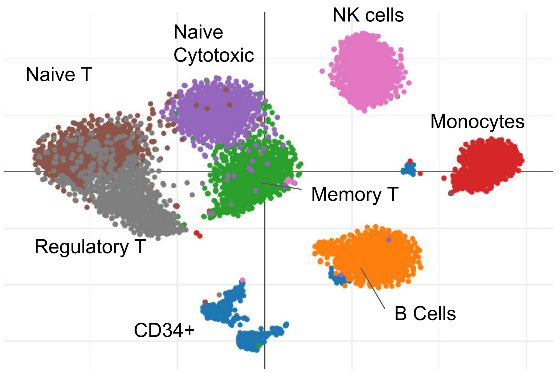


**b) Top cell types for T-Lymphocyte clusters**

| Top cell types for cluster 0 |
|---|
| T-Lymphocytes |
| Lymphocytes |
| CD4-Positive T-Lymphocytes |

| Top cell types for cluster 5 |
|---|
| T-Lymphocytes |
| Lymphocytes |
| CD8-Positive T-Lymphocytes |

| Top cell types for cluster 7 |
|---|
| T-Lymphocytes |
| CD4-Positive T-Lymphocytes |
| T-Lymphocytes, Regulatory |

| Top cell types for cluster 6 |
|---|
| T-Lymphocytes |
| CD8-Positive T-Lymphocytes |
| CD4-Positive T-Lymphocytes |

Figure 5: **a)** tSNE scatterplot showing ground truth labels. **b)** Top 3 most likely cell types as identified by CellMeSH for clusters from Figure 5e.

### 6 Further analysis of the SPLiT-seq spinal cord dataset

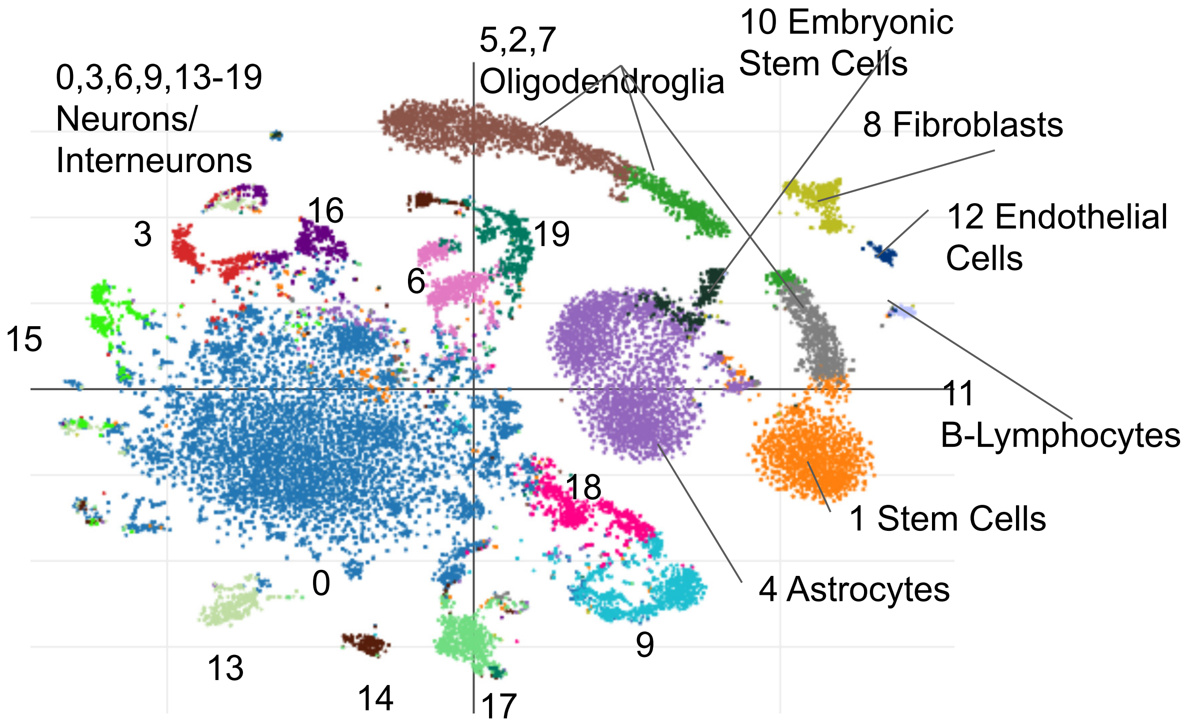


Figure 6: Scatterplot view of the split-seq dataset after further splitting out the neuronal clusters. The non-neuronal clusters are the same as in Figure 6c.

| Cluster | CellMeSH Cell type #1 | Cell type #2 | Cell type #3 | Most common published label |
|---|---|---|---|---|
| 0 | Neurons | Interneurons | Amacrine Cells | Misc Neuron |
| 1 | Stem Cells | Neural Stem Cells | Neuroglia | Oligodendrocyte precursor cell |
| 2 | Oligodendroglia | Schwann Cells | Neurons | Oligodendrocyte |
| 3 | Interneurons | Motor Neurons | Neurons | Inhibitory neuron |
| 4 | Astrocytes | Neuroglia | Neural Stem Cells | Astrocyte |
| 5 | Oligodendroglia | Nerve Fibers, Myelinated | Schwann Cells | Oligodendrocyte |
| 6 | Interneurons | Neurons | Posterior Horn Cells | Excitatory neuron |
| 7 | Oligodendroglia | Stem Cells | Neurons | Oligodendrocyte precursor cell |
| 8 | Stromal Cells | Fibroblasts | Mesenchymal Stem Cells | VLMC (Vascular and leptomeningeal cell) |
| 9 | Neurons | Interneurons | Dopaminergic Neurons | Excitatory neuron |
| 10 | Embryonic Stem Cells | Olfactory Receptor Neurons | Neurons, Afferent | Ependymal cell |
| 11 | B-Lymphocytes | T-Lymphocytes | Bone Marrow Cells | Microglia |
| 12 | Endothelial Cells | Human Umbilical Vein Endothelial Cells | Pericytes | Endothelial cells |
| 13 | Interneurons | Neurons | Purkinje Cells | Inhibitory neuron |
| 14 | Neurons | Posterior Horn Cells | Interneurons | Excitatory neuron |
| 15 | Interneurons | Posterior Horn Cells | Neurons | Inhibitory neuron |
| 16 | Interneurons | Neurons | Stem Cells | Inhibitory neuron |
| 17 | Posterior Horn Cells | Neurons | Interneurons | Excitatory neuron |
| 18 | Neurons | Dopaminergic Neurons | Neurons, Afferent | Excitatory neuron |
| 19 | Posterior Horn Cells | Interneurons | Neurons | Excitatory neuron |

Figure 7: This table shows the top CellMeSH cluster labels for the clusters shown in Figure 3, and the label from the original publication on the right. For most cell types, the top CellMeSH labels belong to the same general category as the published label. Most cells assigned as "Interneurons" were originally labeled as "Inhibitory neurons", while most cells assigned as just "Neurons" or "Posterior Horn Cells" were originally labeled as "Excitatory neurons". The CellMeSH label for cluster 10 is the only one that appears to be clearly incorrect.

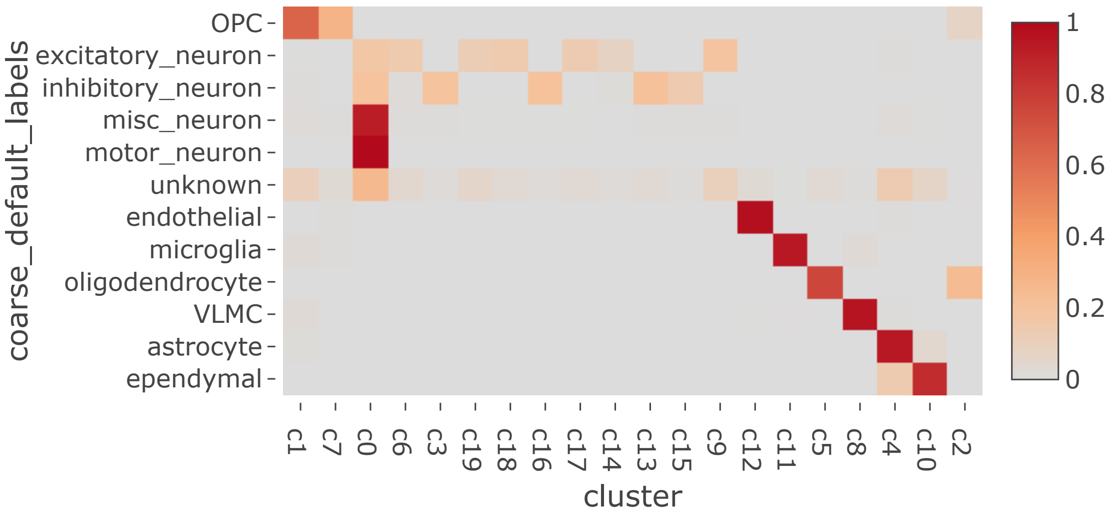


Figure 8: Heatmap showing the (coarse-grained) ground-truth cell types on the y-axis and the UNCURL-App clusters on the x-axis.

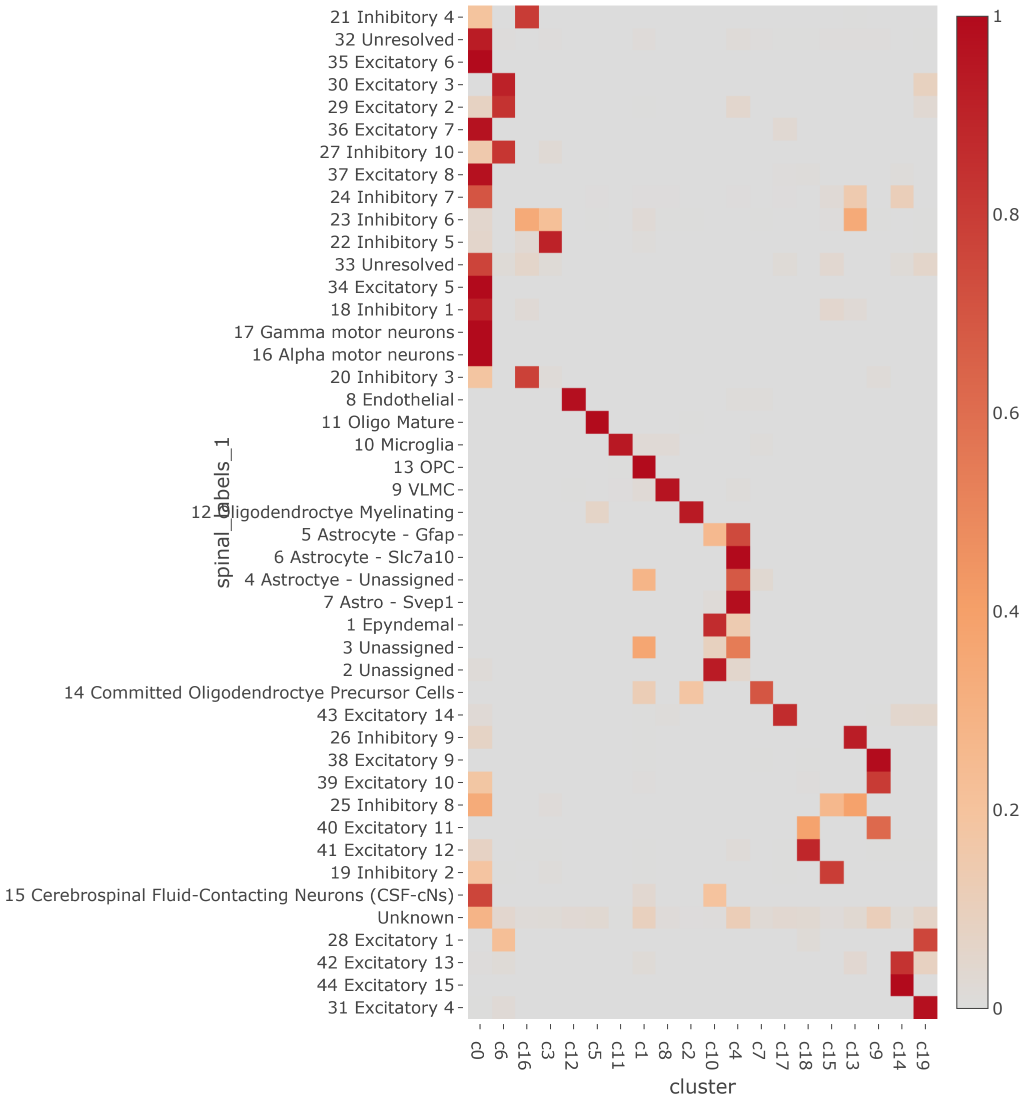


Figure 9: Heatmap showing the (fine-grained) ground-truth cell types on the y-axis and the UNCURL-App clusters on the x-axis. The fine-grained ground-truth labels were the labels provided in the original paper, without combining labels representing similar cell types.

[2] Chris Kang. mnnpy/mnn.py at master · chriscainx/mnnpy.

[3] Sumit Mukherjee, Yue Zhang, Joshua Fan, Georg Seelig, and Sreeram Kannan. Scalable preprocessing for sparse scRNA-seq data exploiting prior knowledge. *Bioinformatics*, 34(13):i124–i132, July 2018.
